# Lifetime genealogical divergence within plants leads to epigenetic mosaicism in the long-lived shrub *Lavandula latifolia* (Lamiaceae)

**DOI:** 10.1101/2020.12.11.421594

**Authors:** Carlos M. Herrera, Pilar Bazaga, Ricardo Pérez, Conchita Alonso

## Abstract

- Epigenetic mosaicism is a possible source of within-plant phenotypic heterogeneity, yet its frequency and developmental origin remain unexplored. This study examines whether the extant epigenetic heterogeneity within long-lived *Lavandula latifolia* (Lamiaceae) shrubs reflects recent epigenetic modifications experienced independently by different plant parts or, alternatively, it is the cumulative outcome of a steady lifetime process.
- Leaf samples from different architectural modules were collected from three *L. latifolia* plants and characterized epigenetically by global DNA cytosine methylation and methylation state of methylation-sensitive amplified fragment length polymorphism markers (MS-AFLP). Epigenetic characteristics of modules were then assembled with information on the branching history of plants. Methods borrowed from phylogenetic research were used to assess genealogical signal of extant epigenetic variation and reconstruct within-plant genealogical trajectory of epigenetic traits.
- Plants were epigenetically heterogeneous, as shown by differences among modules in global DNA methylation and variation in the methylation states of 6-8% of MS-AFLP markers. All epigenetic features exhibited significant genealogical signal within plants. Events of epigenetic divergence occurred throughout the lifespan of individuals and were subsequently propagated by branch divisions.
- Internal epigenetic diversification of *L. latifolia* individuals took place steadily during their development, a process which eventually led to persistent epigenetic mosaicism.

## Introduction

Within-plant variance in phenotypic traits of reiterated, homologous organs that perform the same function (leaves, flowers, fruits, seeds) is often very high, sometimes contributing more to total population-wide variance than differences among individuals (Herrera, 2009, 2017; Palacio *et al*., 2019). Depending on its magnitude and spatio-temporal patterning, this subindividual phenotypic variance can have multiple ecological effects. These include optimizing the exploitation of limiting resources such as light, water or nitrogen (Osada *et al*., 2014; Ponce-Bautista *et al*., 2017; Mediavilla *et al*., 2019), altering the outcome of interactions with animals (Sobral *et al*., 2013, 2014; Shimada *et al*., 2015; Wetzel *et al*., 2016), driving selection on reproductive traits (Austen *et al*., 2015; Dai *et al*., 2016; Arceo-Gómez *et al*., 2017; Kulbaba *et al*., 2017), and enhancing tolerance of environmental unpredictability (Tíscar Oliver & Lucas Borja, 2010; Hidalgo *et al*., 2016). Because of these ecological effects, subindividual variability can eventually influence the fitness of individuals and become itself a target for natural selection, since plants not only have characteristic trait means but also characteristic trait variances and spatio-temporal patterns of subindividual heterogeneity (Herrera, 2009, 2017; Kulbaba *et al*., 2017; Harder *et al*., 2019).

The evolutionary significance of within-plant heterogeneity in phenotypic traits of reiterated structures will depend on its causal mechanisms (Herrera, 2017). Position in relation to external environmental gradients (light, air temperature) or internal developmental axes (nodal position on branches) are factors ordinarily contributing to within-plant variation in phenotypic traits (Herrera, 2009). Subindividual polymorphisms in chromosome number (aneusomaty; D’Amato, 1997) or the DNA sequence of nuclear (Wang *et al*., 2019; Orr *et al*., 2020) and plastid genomes (García *et al*., 2004; Sun *et al*., 2019) can also account for subindividual phenotypic heterogeneity (Whitham *et al*., 1984). The phenotypic effects of these polymorphisms, however, have been investigated on few occasions. Genetic heterogeneity caused by somatic mutations is unlikely to be a pervasive driver of within-plant heterogeneity in wild plants, given the paucity of well-documented genetic mosaics and the extremely low somatic mutation rates reported whenever such mosaics have been found (Padovan *et al*., 2013; Ranade *et al*., 2015; Schmid-Siegert *et al*., 2017; Gerber, 2018; Wang *et al*., 2019; Orr *et al*., 2020). Somatic mutations altering nuclear or plastid DNA sequences are not, however, the only molecular mechanism with a capacity to produce genomic heterogeneity and induce phenotypic variation within individual plants. Cytosine methylation is a major epigenetic mechanism in plants with roles in gene expression, transposon activity, and plant growth and development (Finnegan *et al*., 2000; Cokus *et al*., 2008; Lister *et al*., 2008), hence subindividual heterogeneity in pattern and level of DNA methylation could also partly account for within-plant variation in organ traits (Herrera & Bazaga, 2013; Alonso *et al*., 2018; Herrera *et al*., 2019).

Epigenetic mosaics in which homologous organs in different parts of the same genetic individual differ in extent and/or patterns of DNA methylation have been documented for clonal and non-clonal plants (Bitonti *et al*., 1996; Gao *et al*., 2010; Bian *et al*., 2013; Spens & Douhovnikoff, 2016), and associations between phenotypic heterogeneity and subindividual epigenetic variation, either natural or experimentally induced, have been also found (Herrera & Bazaga, 2013; Alonso *et al*., 2018; Herrera *et al*., 2019). In adult *Lavandula latifolia* (Lamiaceae) plants there is substantial subindividual heterogeneity in global DNA cytosine methylation. Such variation is correlated with within-plant variation in the number and size of seeds produced (Alonso *et al*., 2018), which supports a causal link between epigenetic mosaicism and subindividual phenotypic heterogeneity. In the perennial herb *Helleborus foetidus* (Ranunculaceae), artificial augmentation of within-plant heterogeneity in global DNA cytosine methylation enhanced within-plant variance in phenotypic traits, as predicted by the hypothesis that epigenetic mosaicism can contribute to within-plant variation (Herrera *et al*., 2019).

Two different mechanisms could lead to within-plant epigenetic mosaics and associated phenotypic heterogeneity of the sort documented by Alonso *et al*. (2018) for *L. latifolia*. These mosaics could mostly reflect ephemeral epigenetic modifications experienced recently by different plant parts independently of each other or, alternatively, represent relatively stable somatic conditions reflecting past epigenetic changes which took place at different moments in the plant’s ontogeny. Under this latter scenario, subindividual epigenetic heterogeneity at a given moment in a plant’s lifetime should be genealogically structured, representing the signature of past localized changes within the plant that were propagated and maintained through successive divisions of terminal meristems. This mechanism for internal epigenetic divergence by propagation of stable epimutations is essentially identical to that proposed previously to account for stable subindividual genetic mosaicism via propagation of somatic mutations (Whitham & Slobodchikoff, 1981; Buss, 1983a, b; Whitham *et al*., 1984; Gill *et al*., 1995; Schmid-Siegert *et al*., 2017; Orr *et al*., 2020).

This study evaluates the hypothesis that extant epigenetic heterogeneity within old plants of the evergreen shrub *L. latifolia* is genealogically structured and can thus be considered the outcome of a lifetime process of cumulative epigenetic diversification taking place within plants. Leaf samples from many distinct modules from the same individual plant were characterized epigenetically by measuring global DNA cytosine methylation and assessing the methylation state of a large number of methylation-sensitive anonymous DNA markers. Analyses were accomplished for three wild plants. By combining these data with detailed information on the branching history of each plant, and then applying quantitative analytical methods borrowed from phylogenetic research to assess genealogical signal and reconstruct changes in epigenetic traits, we aim to assess whether extant epigenetic heterogeneity among the modules of the same individual attests their past genealogical trajectories within the plant.

## Materials and methods

### Study species and field methods

*Lavandula latifolia* Med. is a dome-shaped, long-lived evergreen shrub inhabiting the undergrowth of mid-elevation woodlands in the eastern Iberian Peninsula. Branching is dichasial and generally conforms to Leeuwenberg’s development model (Hallé *et al*., 1978; Hallé, 1986). This branching pattern leads to crowns of adult plants being made up of morphological units consisting of distinct leaf clusters borne by short stems, many of which produce one terminal inflorescence in early summer (Alonso *et al*., 2018: Fig. 1). Each of these clusters of even-aged leaves borne by short terminal branchlets will be hereafter termed a ‘module’ following Hallé’s (1986) definition (‘the leafy axis in which the entire sequence of aerial differentiation is carried out, from the initiation of the meristem that builds up the axis to the sexual differentiation of its apex’). Mean seed mass and total seed production vary widely among modules of the same plant (Herrera, 1991, 2000), and this variation was shown by Alonso *et al*. (2018) to be related to subindividual mosaicism in global DNA cytosine methylation. Further details on the natural history, reproductive biology and demography of *L. latifolia* can be found in Herrera (1991), Herrera & Jovani (2010), Herrera & Bazaga (2016), and references therein.

**Fig. 1.**
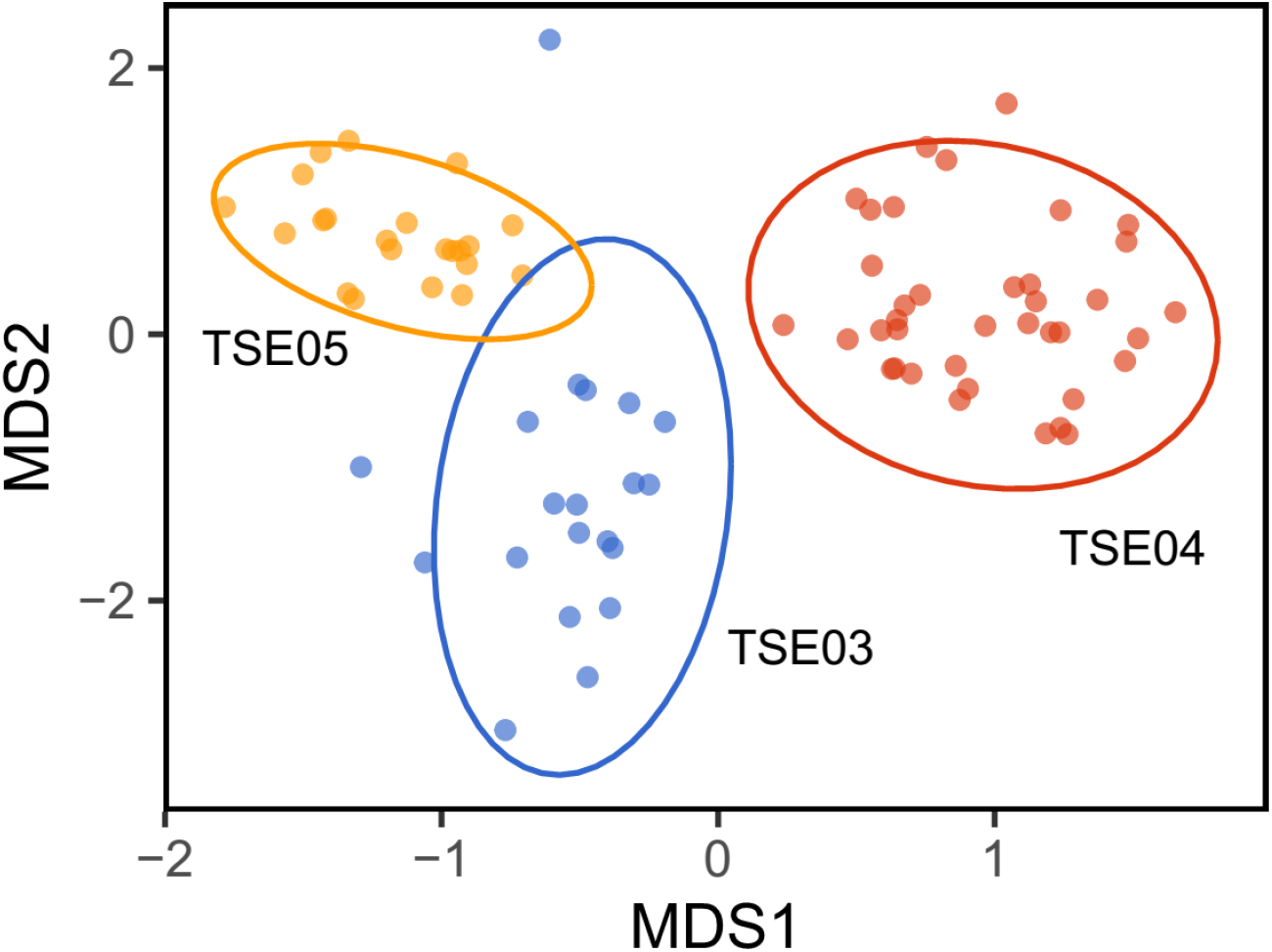
Scatterplot of the *N* = 80 modules (dots) sampled from three *Lavandula latifolia* plants (TSE03, TSE04, TSE05) on the plane defined by nonmetric multidimensional scaling of the matrix of pairwise epigenetic distances (MDS1 and MDS2; coordinates scaled to standard deviation unit and centered to the mean). Epigenetic distances between modules were obtained from the binary matrix of methylation states for the *N* = 400 informative MS-AFLP loci shared by all plants. Ellipses denote the 95% bivariate confidence intervals around individual plant means. A small amount of random variation was added to the location of each point to reveal modules with identical coordinates.

Field sampling for this study was conducted on September 2017 at a large *L. latifolia* population growing near Arroyo Aguaderillos in the Sierra de Cazorla (Jaén province, southeastern Spain). Three of the shrubs whose leaves and seeds had been previously sampled by Alonso *et al*. (2018; plants TSE03, TSE04 and TSE05) were harvested by digging up their roots and brought to the laboratory. Each plant was an individual arising from a single taproot. Fresh leaf samples were collected from as many modules as possible of each plant within a few hours of harvest, subject to the contraint that all leaves in the sampled module should be healthy and free of any visible damage. Samples were placed in paper envelopes, quickly dried at ambient temperature in containers with abundant silica gel, and stored dry at ambient temperature for subsequent DNA extraction. A total of 20, 38 and 22 leaf samples were obtained from as many modules of plants TSE03, TSE04 and TSE05, respectively.

### Laboratory methods

Harvested plants were 23 (TSE03), 29 (TSE04) and 27 (TSE05) years old at time of collection as determined by ring counting (Herrera, 1991). A detailed map of the branching architecture of each plant was drawn by hand, and tips of branchlets from which leaf samples had been collected, and all internal branching nodes leading to them, were tagged and numbered. The linear distances between successive branching nodes, and between the last branching nodes and tips, were measured along branches. The age of every branching node and terminal branchlet (module) was determined by ring counting.

Dried leaf material was homogenized to a fine powder using a Retsch MM 200 mill and total genomic DNA was extracted from approximately 35 mg of ground leaf material using the Bioline ISOLATE II Plant DNA Kit and the manufacturer protocol. Aliquots from the DNA extract of each module (*N* = 80) were used for estimating genome-wide methylation of DNA cytosines and obtaining epigenetic fingerprints as detailed below.

Genome-wide percent cytosine methylation was determined for the leaves of each module using the chromatographic technique described by Alonso *et al*. (2016; see also Alonso *et al*., 2018). Genomic DNA was digested with DNA Degradase PlusTM (Zymo Research, Irvine, CA), a nuclease mix that degrades DNA to its individual nucleoside components. Digested samples were stored at −20°C until analyzed. Two independent technical replicates of DNA hydrolyzate were prepared for each module, and the 160 samples (80 modules x 2 replicates) were processed in randomized order. DNA cytosine methylation was determined for each sample by reversed phase HPLC with spectrofluorimetric detection. Global cytosine methylation was estimated as 100 x 5mdC/(5mdC + dC), where 5mdC and dC are the integrated areas under the peaks for 5-methyl-2’-deoxycytidine and 2’-deoxycytidine, respectively. The position of each nucleoside was determined using commercially available standards (Sigma Aldrich).

Variation among modules of the same plant in epigenetic fingerprint was investigated using a variant of the amplified fragment-length polymorphisms technique (AFLP) which allowed to identify instances of within-plant polymorphism in the methylation state of methylation-susceptible anonymous 5’-CCGG sequences. As we were interested in detecting heterogeneity in genomic DNA methylation profiles among modules from the same genotype, our AFLP method used exclusively primer combinations based on the methylation-sensitive HpaII enzyme. HpaII cleaves 5’-CCGG sequences but is inactive when either or both cytosines are fully methylated, and cleaving may be impaired or blocked when one or both of the cytosines are hemi-methylated (McClelland *et al*., 1994; Roberts *et al*., 2007). In absence of DNA sequence variation among samples, as expected for leaves from the same plant, any within-plant polymorphism in these methylation-sensitive AFLP markers (MS-AFLP hereafter) will reflect subindividual heterogeneity in the methylation state of the associated 5’-CCGG site (see Verhoeven *et al*., 2010; Herrera *et al*., 2012, Herrera & Bazaga, 2013; for applications of this simplified AFLP method in epigenetic studies of plants and fungi). MS-AFLP analyses and fragment scoring were performed following the protocols described in Herrera & Bazaga (2013, 2016). Leaf samples were fingerprinted using eight primer combinations, each with two (HpaII) or three (MseI) selective nucleotides, which were chosen on the basis of repeatability and ease of scoring (Supporting Information Table S1). Scoring error rates were determined for each individual marker by running replicated analyses for 25 leaf samples (31.2% of total), and estimated as the ratio of the number of discordant scores in the two analyses to the total number of replicated samples. To minimize the possibility of spurious within-plant polymorphisms arising from scoring errors, only the *N* = 467 markers with scoring error rates equal to zero were retained for the analyses (Supporting Information Table S1).

### Data analysis

#### Extant epigenetic heterogeneity

All statistical analyses reported in this paper were carried out using the R environment (R Core Team, 2020). Heterogeneity in global cytosine methylation among sampled plants, and among modules of the same plant, was tested by fitting a linear model to the data, treating plants and modules nested within plants as fixed-effect predictors. The contribution of differences between plants, and between modules within plants, to total sample variance in global cytosine methylation were estimated by fitting an intercept-only random effect model to the data using the lmer function of the lme4 package (Bates *et al*., 2015), with plants and modules as hierarchically nested random effects. Confidence intervals of variance estimates were computed using the function confint.merMod in lme4.

For each plant, a module x MS-AFLP marker binary matrix was obtained whose elements were the methylation state of each marker in the given module (1 = unmethylated; 0 = methylated). In each matrix, only those markers occurring unmethylated in at least one module of the plant were retained for analysis (‘informative markers’ hereafter), because our MS-AFLP procedure did not allow to discriminate between homogeneous methylation and fragment absence for those markers which did not occur in any module of a plant. Another binary matrix was obtained for all plants and modules combined using only those markers that were simultaneously informative in all plants (*N* = 400). Multivariate analyses of within-and among-plant variation in epigenetic fingerprints were then conducted on these data, which included nonmetric multidimensional scaling (function metaMDS in package vegan; Oksanen *et al*., 2019) and analysis of molecular variance (adonis function in vegan) on the pairwise matrix of Jaccard dissimilarity between modules.

#### Within-plant genealogy of epigenetic heterogeneity

Two Newick-formatted genealogical trees were constructed for each plant by collating the topological information from drawings of branching architecture and the quantitative data on linear or temporal distances between branching nodes. In these genealogical trees the modules sampled were at the tips and branch lengths were either the linear distance between branching nodes or their age difference in years (Supporting Information Fig. S1). Although growth- and age-based branch lengths were correlated in the three plants (*R*^2^ = 0.57-0.61), we used trees based on these two different metrics to explore whether epigenetic changes taking place over a plant’s lifetime were best explained in terms of the amount of growth or the time elapsed between successive branching nodes.

The dichotomous branching trees used here to depict genealogical relationships among extant modules of *L. latifolia* plants are conceptually equivalent to the phylogenetic trees commonly used to depict the evolution of contemporary taxa by descent with modification. There is the advantageous difference that our genealogical trees are errorless pedigrees representing true relationships rather than uncertain inferential hypotheses as it usually happens with phylogenetic trees (Felsenstein, 2004). Methods borrowed from phylogenetic research were used to investigate the genealogical component of extant within-plant epigenetic mosaicism in *L. latifolia* shrubs (see Orr *et al*., 2020, for a comparable approach). ‘Genealogical tree’, ‘genealogical character estimation’ and ‘genealogical signal’ will be used hereafter as the within-plant equivalents to ‘phylogenetic tree’, ‘ancestral character estimation’ and ‘phylogenetic signal’, respectively (Paradis, 2012; Münkemüller *et al*., 2012). Assessment of genealogical signal and genealogical character estimation will be consistently used throughout this paper to evaluate the genealogical basis of extant epigenetic mosaicism within *L. latifolia* plants. The first approach provides a quantitative assessment at the whole plant level of the association betweeen trait similarity and proximity in the genealogy, while the second will inform on the spatio-temporal patterns of epigenetic changes taking place within individual plants over their lifetimes.

For continuous traits (global DNA cytosine methylation and coordinates of modules on axes from nonmetric multidimensional scaling), genealogical character estimations were carried out with function contMap from the phytools package (Revell, 2012). Genealogical signal was tested with the philoSignal function in the philosignal package (Keck et al., 2016) and Moran’s *I* method, which relies on an autocorrelation approach, makes no assumptions on model of change and incorporates information on branch length (Münkemüller *et al*., 2012). Within-plant genealogical signal in the methylation state of individual MS-AFLP markers (0 = methylated, 1 = unmethylated) was tested using Fritz & Purvis’ (2010) *D* statistic for binary traits, a measurement of character dispersion on the genealogy. In these analyses, only the most informative polymorphic markers (frequency of commonest methylation state < 0.85) were considered for each plant (*N* = 3, 6 and 6 markers for plants TSE03, TSE04 and TSE05, respectively). Computations were conducted with function phylo.d from the caper package (Orme *et al*., 2018). Randomization tests were used to assess statistical significance of differences between observed *D* values and expectations from random (*D* = 1) and Brownian motion (*D* = 0) distributions of methylation state of individual markers across tips of genealogical trees (Fritz & Purvis, 2010). Transition rates between the methylated and unmethylated states of individual markers on plant genealogies were explored by fitting ‘Equal rates’ (ER) and ‘All rates different’ (ARD) discrete evolution models to genealogical trees using the fitDiscrete function in the geiger package (Pennell *et al*., 2014). Akaike information criterion (AIC) for fitted ER and ARD models were compared with the aic.w function of the phytools package. The stochastic mapping procedure of Bollback (2006), as implemented in function make.simmap of the phytools package, was used for genealogical character estimation of the methylation state of markers with genealogical signal within plants.

## Results

### Extant epigenetic heterogeneity

There was substantial subindividual heterogeneity in global DNA cytosine methylation among the even-aged leaf samples from different modules. In addition to differences among plants (*F*_2,92_ = 12.18, *P* = 0.00002), cytosine methylation also differed among modules of the same plant (*F*_77,92_ = 1.59, *P* = 0.017). Estimated within-plant variance in global cytosine methylation (0.119; 95% confidence interval = 0.015–0.252) was roughly comparable to among-plant variance (0.100; 95% confidence interval = 0.0056–0.647), further stressing the quantitative importance of within-plant variation in the extent of genomic methylation.

Modules from the same plant were broadly scattered on the plane defined by axes from nonmetric multidimensional scaling of the pairwise dissimilarity matrix (Fig. 1), thus revealing substantial within-plant epigenetic heterogeneity in the multivariate space defined by those informative MS-AFLP markers which were shared by all plants (*N* = 400). Analysis of molecular variance on the matrix of pairwise dissimilarities between modules revealed that 29.2% of total epigenetic variance in the sample was accounted for by differences among modules of the same plant.

Subindividual epigenetic heterogeneity was also apparent when informative MS-AFLP markers were considered individually. The three plants studied were closely similar with regard to the number of informative markers (range = 427-431 markers; Table 1). In every plant a small, but non-negligible fraction of these markers (range = 5.8-7.7%; Table 1) were polymorphic in methylation state among modules of the same plant, i.e., occurred in the methylated and unmethylated states in different parts of the same shrub. About one third of polymorphic informative markers occurred predominantly in the methylated state, and two thirds occurred predominantly in the unmethylated state (Table 1). About 20% of subindividually polymorphic loci (*N* = 75, all plants combined) were polymorphic in more than one plant (Supporting Information Fig. S2).

**Table 1.** Variation among modules of the same plant in methylation state of MS-AFLP markers. See supporting Information Fig. S2 for distribution among plants of shared and unique polymorphic markers.

| Plant | Modules sampled | Informative markers* | Subindividually polymorphic markers |  |  |
| --- | --- | --- | --- | --- | --- |
|  |  |  | Total | Predominant state: |  |
|  |  |  |  | Methylated | Unmethylated |
| TSE03 | 20 | 431 | 33 | 11 | 22 |
| TSE04 | 38 | 427 | 33 | 9 | 24 |
| TSE05 | 22 | 427 | 25 | 8 | 17 |

\* Out of a total of the $N = 467$ markers considered whose estimated scoring error rates were equal to zero. Individual markers were considered informative for a given plant if they were recorded in at least one of its modules (see Materials and Methods: Data analysis).

### Within-plant genealogy of epigenetic heterogeneity

Within-plant heterogeneity in global DNA cytosine methylation had genealogical signatures in two plants (TSE04 and TSE05), as denoted by statistically significant or marginally significant Moran’s *I* autocorrelations, irrespective of the branch length metric used to construct genealogical trees (Table 2). Genealogical character estimations revealed some nodes early in the plants’ lives whose descendant branches were consistently characterized until the time of collection by divergent values of global cytosine methylation (marked by arrows in Fig. 2).

**Fig. 2.**
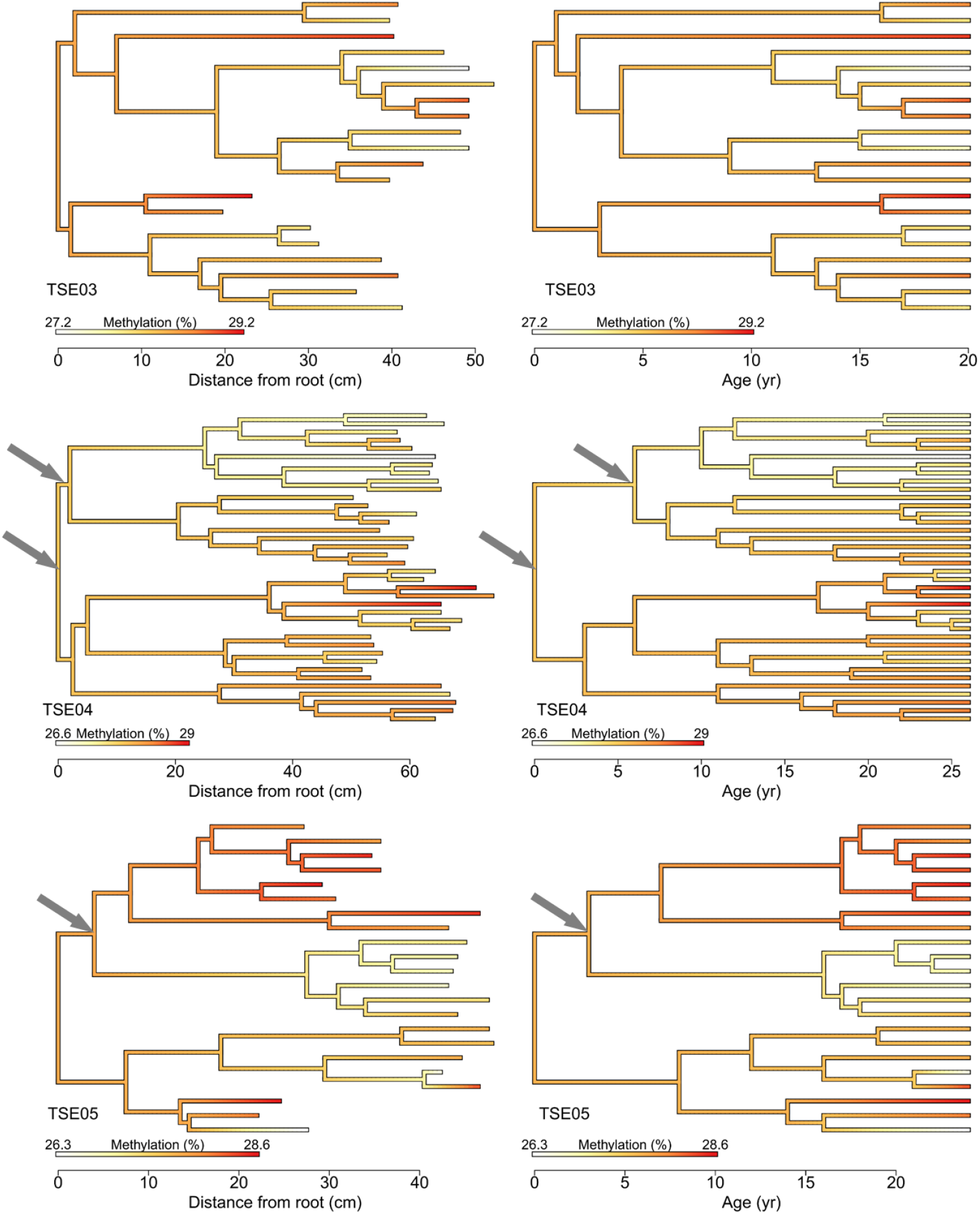
Genealogical character estimation of within-plant changes in global DNA cytosine methylation for the three *Lavandula latifolia* plants studied (TSE03, TSE04, TSE05). Two genealogical trees were used for each plant, whose branch lengths were either linear distances (left column) or age differences (right column) between nodes (Supporting Information Fig. S1). Estimated changes in trait value along branches are color-mapped on each tree according to the scales shown. Limits of color scales differ slightly among plants because they were adjusted in each case to the corresponding minimum and maximum values. The arrows mark branching nodes referred to in the text.

**Table 2.**
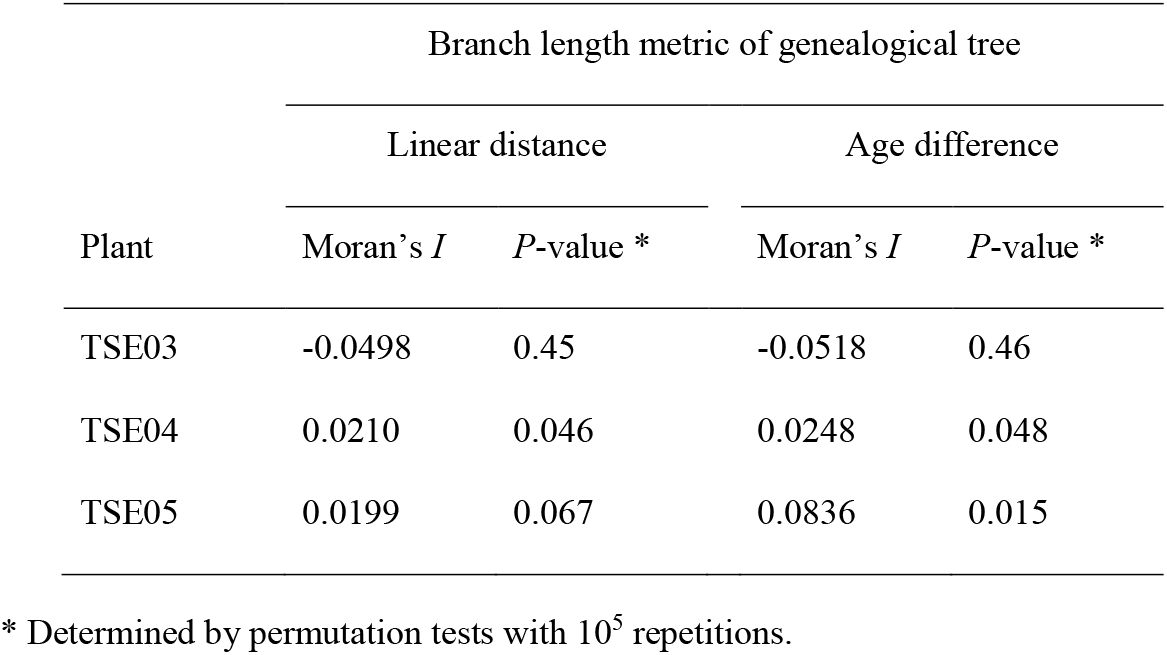
Tests of genealogical signal for among-module variation in global DNA cytosine methylation within the three *Lavandula latifolia* plants studied.

| Plant | Branch length metric of genealogical tree |  |  |  |
| --- | --- | --- | --- | --- |
|  | Linear distance |  | Age difference |  |
|  | Moran's *I* | *P*-value * | Moran's *I* | *P*-value * |
| TSE03 | -0.0498 | 0.45 | -0.0518 | 0.46 |
| TSE04 | 0.0210 | 0.046 | 0.0248 | 0.048 |
| TSE05 | 0.0199 | 0.067 | 0.0836 | 0.015 |

\* Determined by permutation tests with $10^5$ repetitions.

Heterogeneity among modules of the same plant in multilocus epigenetic fingerprints, as assessed by coordinates from nonmetric multidimensional scaling of pairwise distance matrices (MDS1 and MDS2), had statistically significant genealogical signals in plants TSE03 (axis MDS1) and TSE04 (axes MDS1 and MDS2), irrespective of branch length metric used to construct genealogical trees (Table 3). Genealogical character estimations for MDS1 and MDS2 revealed one or more branching nodes early in the lives of the plants studied whose descendant branches were subsequently characterized by divergent multilocus epigenetic fingerprints until the time of collection (nodes marked by arrows, Fig. 3).

**Fig. 3.**
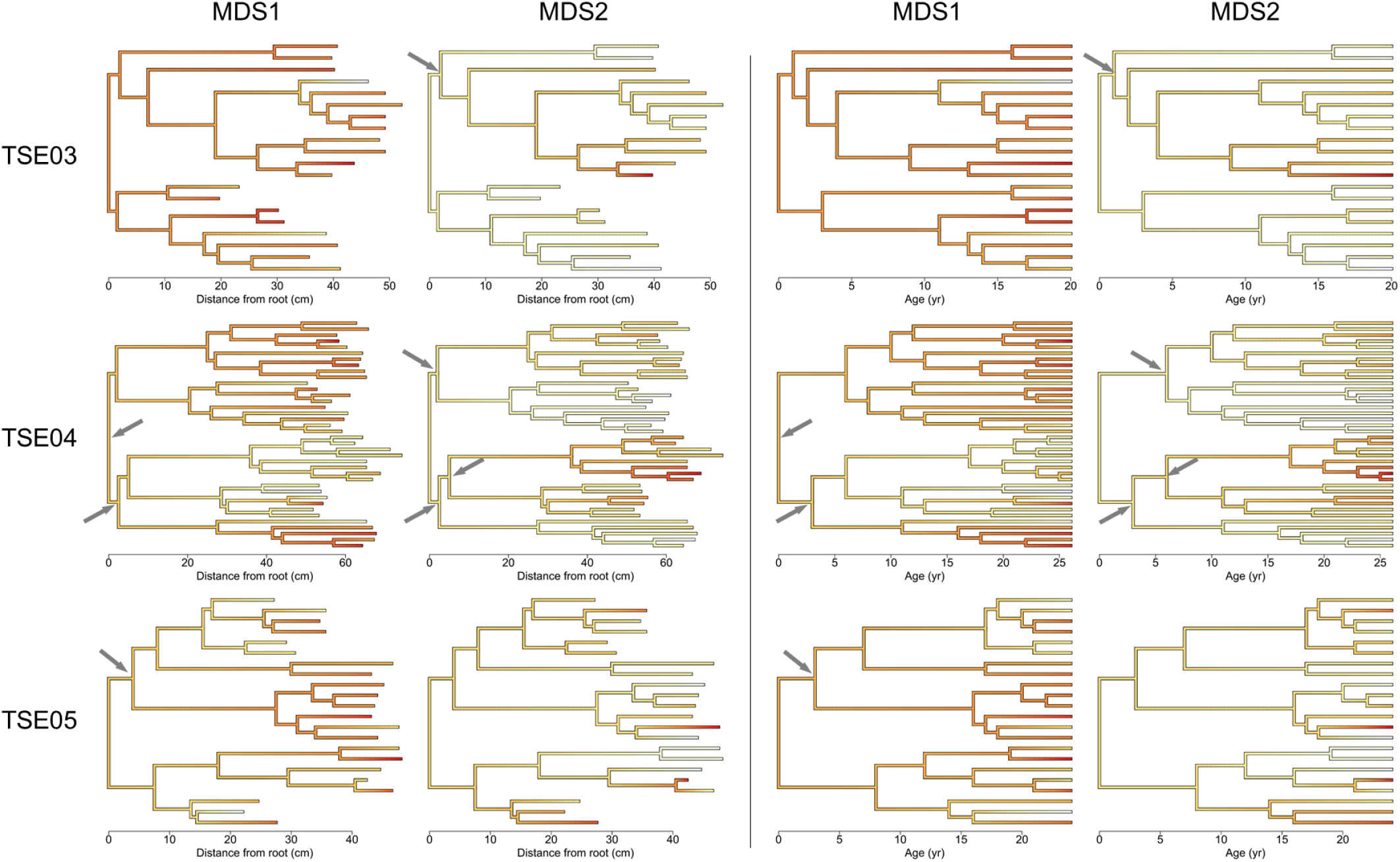
Genealogical character estimation within individual *Lavandula latifolia* plants of changes in multilocus epigenetic fingerprints of individual modules, as described by their coordinates on the axes obtained from nonmetric multidimensional scaling of pairwise distance matrices (MDS1 and MDS2). Two trees were used for each plant, whose branch lengths were either linear distances (two left columns) or age differences (two right columns) between nodes. Estimated changes in trait value along branches are color-mapped on each tree. In each tree, the color scale was defined by the minimum (white) and maximum (red) values for the corresponding plant and axis (see Fig. 1). Scales have been omitted to reduce cluttering. The arrows mark branching nodes referred to in the text.

**Table 3.** Tests of genealogical signal for among-module variation in multilocus epigenetic fingerprints, assessed by coordinates from nonmetric multidimensional scaling (Fig. 1).

| Plant | Coordinate | Branch length metric of genealogical tree |  |  |  |
| --- | --- | --- | --- | --- | --- |
|  |  | Linear distance |  | Age difference |  |
|  |  | Moran's *I* | *P*-value * | Moran's *I* | *P*-value * |
| TSE03 | MDS1 | -0.0529 | 0.46 | -0.0610 | 0.53 |
|  | MDS2 | 0.0437 | 0.0074 | 0.0440 | 0.010 |
| TSE04 | MDS1 | 0.0513 | 0.0055 | 0.0673 | 0.0039 |
|  | MDS2 | 0.1452 | <10<sup>-5</sup> | 0.2070 | <10<sup>-5</sup> |
| TSE05 | MDS1 | -0.0576 | 0.56 | -0.0449 | 0.43 |
|  | MDS2 | -0.0894 | 0.86 | -0.0871 | 0.79 |

\* Determined by permutation tests with 10<sup>5</sup> repetitions.

Variation across modules of the same plant in the methylation state of the most polymorphic MS-AFLP markers was significantly related to module genealogy (Fritz-Purvis’ *D* < 1) in about two thirds of instances tested (15 markers x 2 branch length metric combinations) (Fig. 4; Supporting Information Table S2). One, five and three markers exhibited genealogical signal in plants TSR03, TSE04 and TSE05, respectively (Fig. 4). The *D* statistic did not depart significantly from Brownian motion expectations (*D* = 0) in any of these cases (Fig. 4). Genealogical character estimations of within-plant variation in methylation state for markers with genealogical signal are shown in Fig. 5 on trees whose branch lengths are age differences (results were closely similar for trees based on linear distances between nodes; Supporting Information Fig. S3). Genealogical clumping of marker methylation state at tree tips (modules) was evident in all cases, although the size of clumps and their time of divergence along the plant’s lifetime ranged widely among markers. In some cases the initial divergence in methylation state occurred when plants were ≤ 5 yr old, and its subsequent propagation over many years of branching without methylation change eventually formed large genealogical clumps (e.g., TG_CTA_297, TC_CCT_367). In other cases, in contrast, the initial methylation divergence at the base of a clump occurred when plants were already ≥ 15 yr old, and the resulting genealogical clumps were smaller and involved fewer modules (TC_CGC_347, TC_CCT_200; Fig. 5). Some instances of recent reversals in methylation state were apparent within the genealogically oldest clumps (e.g., TG_CTA_297, TC_CCT_367; Fig. 5).

**Fig. 4.**
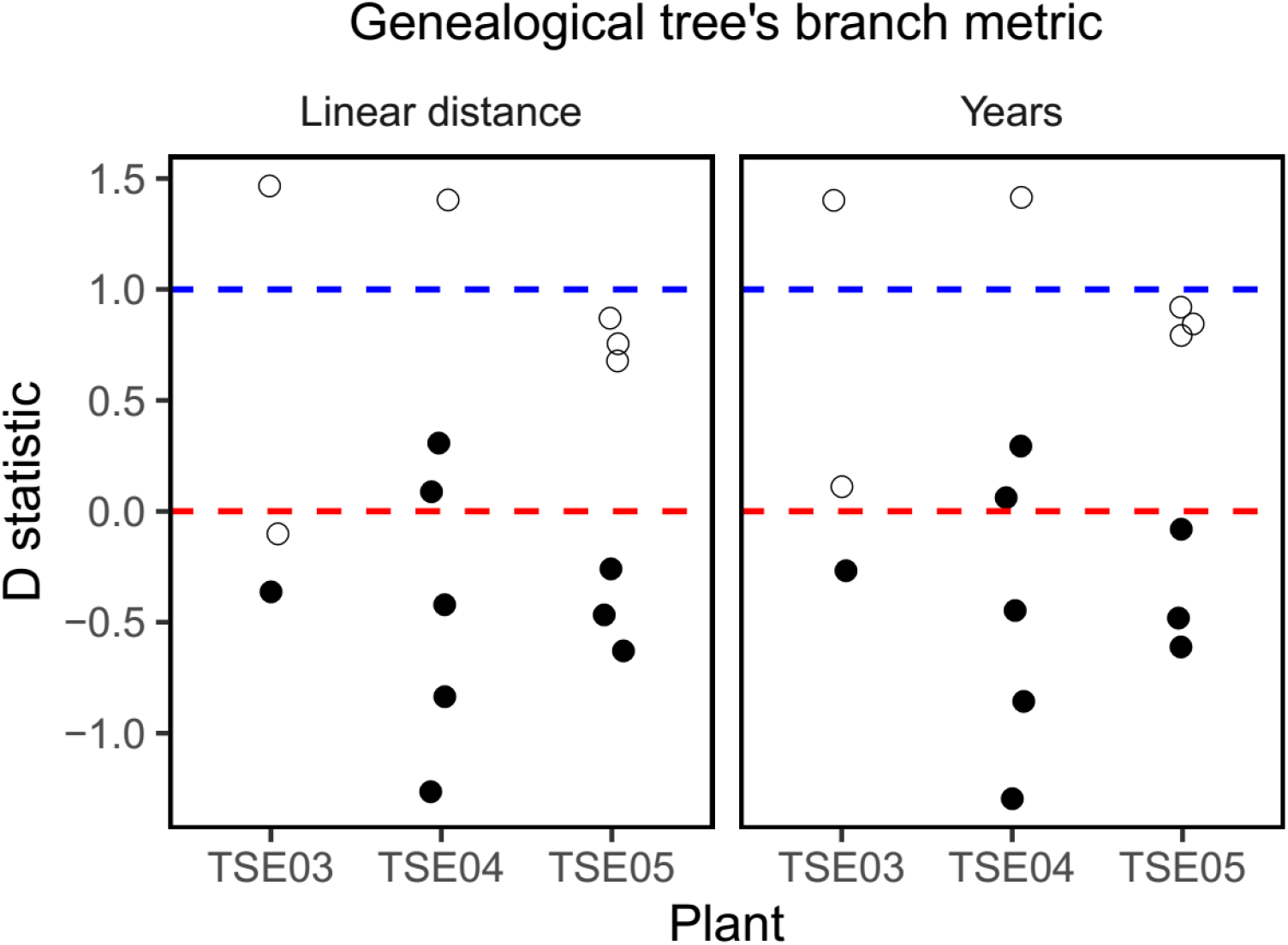
Summary of analyses of within-plant genealogical signal (Fritz-Purvis’ *D* statistic) in the methylation state of highly polymorphic MS-AFLP markers (frequency of commonest methylation state < 0.85; *N* = 3, 6 and 6 markers for plants TSE03, TSE04 and TSE05, respectively). Blue and red dashed lines mark expected values from random and Brownian motion distributions of methylation state across tips of genealogical trees. Filled dots denote markers whose methylation state simultaneously exhibited significant genealogical clumping within plants (*D* significantly < 1) and nonsignificant departure from Brownian motion expectation (*D* = 0). See Supporting Information Table S2 for detailed numerical results.

**Fig. 5.**
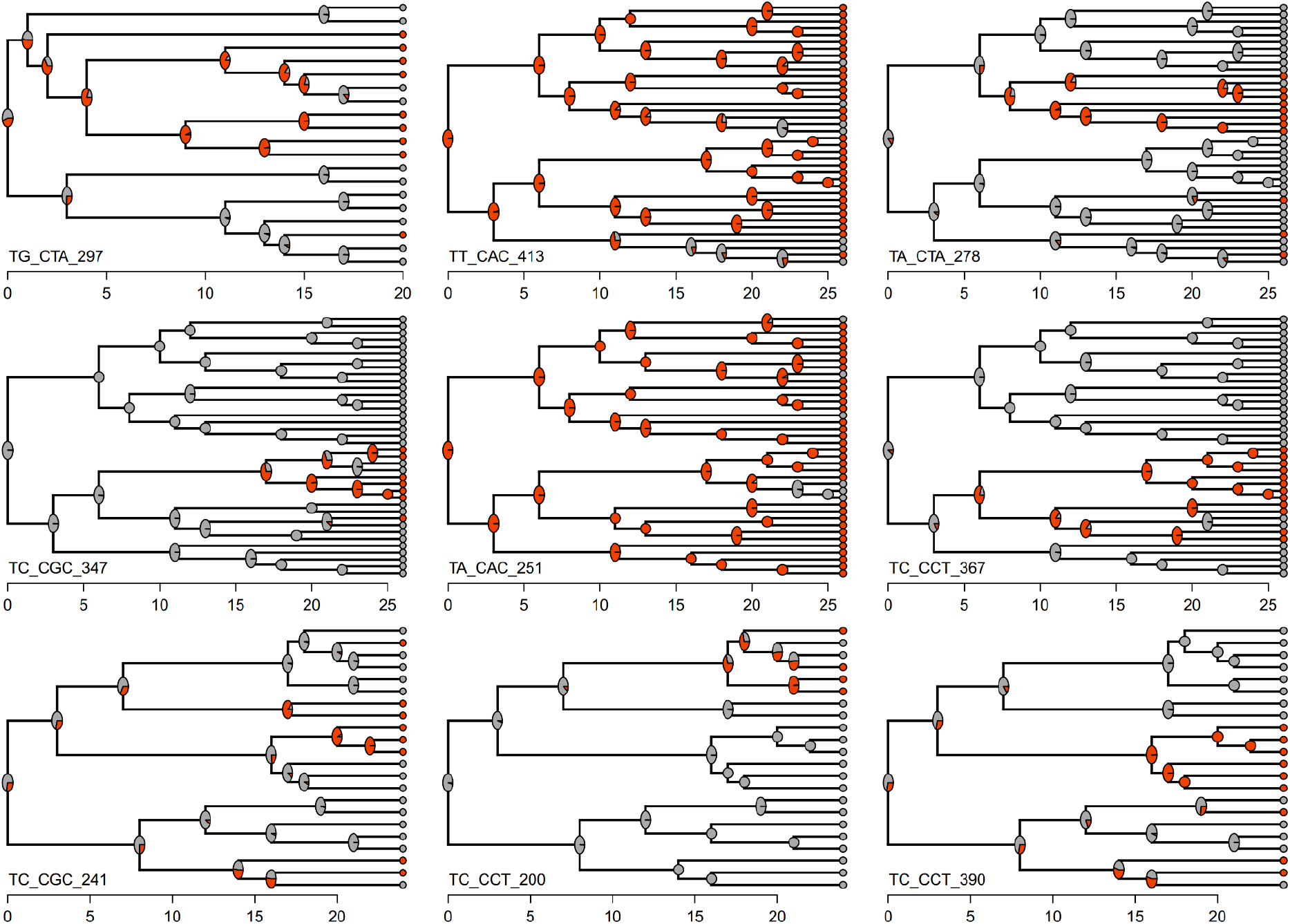
Within-plant genealogical character estimation of the methylation state of nine highly polymorphic MS-AFLP markers with significant genealogical signal (Fig. 4, and Supporting Information Table S2). For each marker, methylation state in the sampled modules (tree tips) and estimated posterior probabilities at nodes are coded as grey (methylated) or orange (unmethylated). Tree branch lengths represent differences in age between nodes and units of horizontal axes are years. Markers are identified by primer combination and fragment size in base pairs (Supporting Information Table S1), and correspond to plants TSE03 (TG_CTA_297), TSE04 (TT_CAC_413, TA_CTA_278, TC_CGC_347, TA_CAC_251, TC_CCT_367) and TSE05 (TC_CGC_241, TC_CCT_200, TC_CCT_390).

Paired ‘Equal rates’ (ER) and ‘All rates different’ (ARD) discrete evolution models fitted to within-plant methylation state data for individual markers generally provided better support for the ER model (63.3% of instances; Supporting Information Table S2). When ARD models provided a better fit (36.7% of instances), mean (± SE) transition rate from the unmethylated to the methylated state (0.360 ± 0.029) was only slightly higher than the transition rate from the methylated to the unmethylated state (0.295 ± 0.022) (Supporting Information Table S2).

## Discussion

### Extant subindividual epigenetic heterogeneity and its origin

Plants of *L. latifolia* were epigenetically heterogeneous at time of collection. Leaves from different modules in the same plant differed in global DNA methylation and MS-AFLP profiles at multilocus and single-marker levels. These findings extend those of Alonso *et al*. (2018) for this species showing within-plant heterogeneity in global methylation for a superset of the plants considered here. Global methylation may vary within species or individuals in relation to plant age or tissue of origin (Mankessi *et al*., 2011; Vining *et al*., 2012; Alonso *et al*., 2017; Gao *et al*., 2019), but none of these factors can account for heterogeneous genomic methylation within plants of *L. latifolia* found here and by Alonso *et al*. (2018), since all DNA samples from the same plant were from even-aged leaf cohorts. The same applies to within-plant heterogeneity in epigenetic fingerprint and methylation state of MS-AFLP markers.

A demographic study on the study population revealed that mean longevity of *L. latifolia* individuals that flowered at least once during their lifetimes was 22 years, and only ~7% of these lived for >30 years (Herrera & Bazaga, 2016). Since plants included in this study were 23-29 yr old at time of collection, our results refer to the older age class in the population. Insofar as within-plant patterning of epigenetic features is a cumulative process taking place over a plant’s lifetime, as suggested by this study, such patterning would possibly have been weaker or harder to detect in younger individuals. This is supported by variation among plants studied in frequency of significant genealogical signals of epigenetic features, which tended to increase from the youngest (TSE03) to the oldest (TSE04) individual. It should also be kept in mind that insufficient statistical power probably hindered detection of genealogical signal in the younger plants. With very small trees (*N* ~ 20 tips, as in TSE03 and TSE05), all methods for detecting genealogical signal have high Type II errors (Blomberg *et al*., 2003; Münkemüller *et al*., 2012). Larger genealogical trees from species with longer longevities (e.g., trees) should be most favorable for the detection of genealogical signal.

Trees used for assessing genealogical signal in extant epigenetic heterogeneity represent the developmental pedigree of modules at the tips, and describe the ontogenetic unfolding of each individual over its lifetime. All modules in a plant derive from the same ancestor, namely the initial seedling arising from a seed, and genealogical trees depict the topology of descendant lineages arising from branching events. Pairs of modules physically closer at tips of a tree are also historically and developmentally closer to their most recent common ancestor module than pairs of modules located farther away in the tree. These relationships, along with the regularly dichasial branching pattern that characterizes *L. latifolia* shrubs, justify our application of methods from phylogenetic research to assess genealogical signal and perform genealogical reconstructions of within-plant epigenetic changes (see also Orr *et al*., 2020). These methods could be used for the same purpose on other woody perennials that follow Leeuwenberg’s model of architecture (see, e.g., Hamilton, 1985; Hallé, 1986; Navarro *et al*., 2009; for tropical and non-tropical examples).

Results of this study agree with expectations from the hypothesis advanced by Alonso *et al*. (2018) that subindividual variation in epigenetic features of *L. latifolia* plants was the consequence of the concerted action of plant sectoriality (plant body’s compartmentalization into physiologically semi-autonomous subunits; Watson, 1986) and the differential action on plant parts of some factor(s) inducing persistent changes in extent and/or patterns of DNA cytosine methylation (e.g., pathogens, herbivores, insolation, UV light, water shortage, nitrogen deficiency; reviewed by Alonso *et al*., 2016). Sectoriality will constrain the horizontal circulation of phloem-mobile molecules that regulate DNA methylation (McGarry & Kragler, 2013; Lewsey *et al*., 2016), thus contributing to maintain within-plant heterogeneity in epigenetic features arising from random epimutations or localized responses to environmental agents, as previously suggested in relation to other subindividually variable traits (Orians & Jones, 2001; Herrera, 2009). We found here that extant within-plant heterogeneity in epigenetic features (global DNA cytosine methylation, MS-AFLP multivariate fingerprint, methylation state of specific MS-AFLP markers) exhibited statistically significant genealogical signals. Results were considerably robust irrespective of whether branch lengths of genealogical trees were linear or temporal distances between nodes. This points to an equivalence of ageing and growing as ultimate agents of within-plant epigenetic diversification over a plant’s lifetime.

Genealogical character reconstructions revealed that early events of internal epigenetic divergence took place when plants were still very young (< 5 yr), before reaching the age of first reproduction (Herrera & Bazaga, 2016). In general, the timing of epigenetic modifications spanned the entire lifespan of individuals, thus revealing that epigenetic features experienced steady changes throughout individual plants’ lives. This was particularly evident in the case of changes in methylation state of subindividually polymorphic MS-AFLP markers, where changes conforming to a Brownian motion model took place over the life of individuals with about similar estimated probabilities in both directions, and even the reversion to the ‘ancestral’ methylation state could be documented. A corollary of this finding is that *L. latifolia* individuals can produce slightly different epigenetic fingerprints over its lifetime if sampled repeatedly over a sufficiently broad timespan. This expectation is upheld by preliminary results for plants from our study population which were sampled on two occasions nine years apart (C. M. Herrera, *unpublished data*).

### Toward an epigenetic mosaicism hypothesis

Modular construction by continual organogenesis and reiterated production of homologous structures is a quintessential plant feature which has motivated the consideration of plant individuals as non-unitary metapopulations of semi-autonomous modules, a notion departing from the common zoocentric definition of organismic individuality (Gerber, 2018). This view led to the incorporation of selection at the subindividual level as a possible evolutionary mechanism (Buss, 1983a, b; Pineda-Krch & Poore, 2004), and provided the foundations for the ‘genetic mosaicism hypothesis’ (GMH) (Whitham & Slobodchikoff, 1981; Whitham *et al*., 1984; Gill *et al*., 1995). The following premises synthesize the GMH (slightly modified from Gill *et al*., 1995): (i) spontaneous mutations occur among the proliferating meristems; (ii) the meristematic and modular basis of plant development assures that many of these mutations are preserved and expanded hierarchically among modules as the plant grows; (iii) the differential growth and survival of ramets, branches and shoots should alter the genotypic configuration of the plant as it grows; and (iv) the within-plant phenotypic heterogeneity arising from genotypic heterogeneity will affect individual fitnesss through effects on the progeny, plant responses to the environment, or responses of animal consumers (Whitham & Slobodchikoff, 1981; Whitham *et al*., 1984; Gill *et al*., 1995; Herrera, 2009).

Studies focusing on subindividual genetic heterogeneity in wild plants have produced few good examples of genetic mosaicism in non-clonal woody plants, and generally documented very low somatic mutation rates (Cloutier *et al*., 2003; Padovan *et al*., 2013; Ranade *et al*., 2015; Schmid-Siegert *et al*., 2017; Wang *et al*., 2019; Orr *et al*., 2020; see also Herrera, 2009, for review). This tends to deny the evolutionary importance of genetic mosaicism advocated by the GMH (Pannell & Eppley, 2004; Gerber, 2018). In contrast, the few investigations that have so far addressed the possibility of subindividual variation in epigenetic features among homologous organs have found relatively high frequencies of somatic epigenetic variants, discernible within-plant epigenetic mosaicism, and/or relationships between subindividual epigenetic heterogeneity and within-plant phenotypic variation (Bitonti *et al*., 1996; Herrera & Bazaga, 2013; Alonso *et al*., 2018; Herrera *et al*., 2019; Hofmeister *et al*., 2020). The present study has extended these previous findings by showing that steady epigenetic diversification over plants’ lifetimes can lie behind extant subindividual epigenetic mosaics.

Taken together, results obtained so far bearing on subindividual epigenetic variation motivate our proposal of an ‘epigenetic mosaicism hypothesis’ (EMH) consisting of exactly the same elements i-iv above as the original GMH but where the terms ‘mutation’, ‘genetic’ and ‘genotype’ are replaced with ‘epimutation’, ‘epigenetic’ and ‘epigenotype’, respectively. Two additional components of GMH, namely inheritance of somatic mutations and impact of mosaicism on individual fitness, will often apply to EMH as well. Transgenerational epigenetic inheritance has been documented for model and non-model plants (Jablonka & Raz, 2009; Hauser *et al*., 2011; Quadrana & Colot, 2016). In *L. latifolia* there is extensive transgenerational transmission of genome-wide global cytosine methylation levels and methylation state of anonymous epigenetic markers (Herrera *et al*., 2018). Although the ecological impact has been rarely investigated, there is also evidence that within-plant epigenetic mosaicism can influence plant fitness (Alonso *et al*., 2018; Herrera *et al*., 2019).

By incorporating the within-plant realm to the already well-accepted consensus that epigenetic variation is an important source of phenotypic variance among individuals and populations (Bossdorf *et al*., 2008, 2010; Lira-Medeiros *et al*., 2010; Medrano *et al*., 2014; Kooke *et al*., 2015; Groot *et al*., 2018), the EMH offers a particularly favorable arena for formulating and testing novel hypotheses on the ecological and evolutionary roles of epigenetic variation while holding constant the influence of genetic factors. For example, lifetime internal epigenetic diversification within individuals, as documented here for *L. latifolia*, may represent a mechanism of ‘exploration’ of the epigenetic landscape endowing each plant with a broader phenotypic space to cope with challenges of the abiotic and biotic environment. The breadth of such epigenetic sampling (i.e., within-individual epigenetic/phenotypic variance) should vary depending on life expectancy and species-specific patterns of meristem divisions related to the architectural model. Simple predictions from hypotheses framed around the EMH are amenable to experimentation by manipulating within-plant epigenetic heterogeneity while keeping genetic background constant, e.g., by localized application of chemical agents which alters methylation and monitoring effects on phenotypic heterogeneity and ecological consequences (Herrera *et al*., 2019). These investigations are bound to contribute new insights on the mechanistic basis and ecological and evolutionary implications of the within-plant component of phenotypic variance in plant populations.

## Acknowledgements

We are grateful to Esmeralda López Perea for laboratory assistance. Consejería de Medio Ambiente, Junta de Andalucía, authorized this research and provided invaluable facilities at the study area. Partial support for this study was provided by grants CGL2016-76605-P (Ministerio de Economía y Competitividad, Spanish Government), PID2019-104365GB-I00 (Ministerio de Ciencia e Innovación, Spanish Government), and P18-FR-4413 (Consejería de Transformación Económica, Industria, Conocimiento y Universidades, Andalusian Government).

## Author contributions

CMH designed the research, did field work, conducted data analyses, and led the writing; CA contributed data analyses and interpretations, and supervised laboratory work; RP and PB performed HPLC and MS-AFLP analyses, respectively; all authors reviewed and edited the manuscript.

## Supporting Information

Additional Supporting Information may be found online in the Supporting Information section at the end of the article.

**Fig. S1** Tree representations of genealogical relationships among the modules sampled in each of the three *Lavandula latifolia* individuals studied.

**Fig S2** Venn diagrams showing the distribution among plants of subindividually polymorphic methylation-sensitive AFLP markers.

**Fig. S3** Within-plant genealogical character estimation of the methylation state of polymorphic MS-AFLP markers on trees whose branch lengths are linear distances between nodes.

**Table S1** Primer combinations and number of fragments that were used in the MS-AFLP analyses of leaf DNA samples.

**Table S2** Summary of tests of within-plant genealogical signal, and fits of discrete character models, for highly polymorphic individual MS-AFLP markers.

