## Supporting Information for "Lifetime genealogical divergence within plants leads to epigenetic mosaicism in the long-lived shrub *Lavandula latifolia* (Lamiaceae)"

**Fig. S1** Tree representations of the genealogical relationships among the modules sampled in each *Lavandula latifolia* plant studied (tree tips). Two trees were obtained for each plant whose edges represented either linear distances (“growth”, left column, in centimeters) or age differences (“age”, right column, in years) between nodes. Edge lengths shown in red.

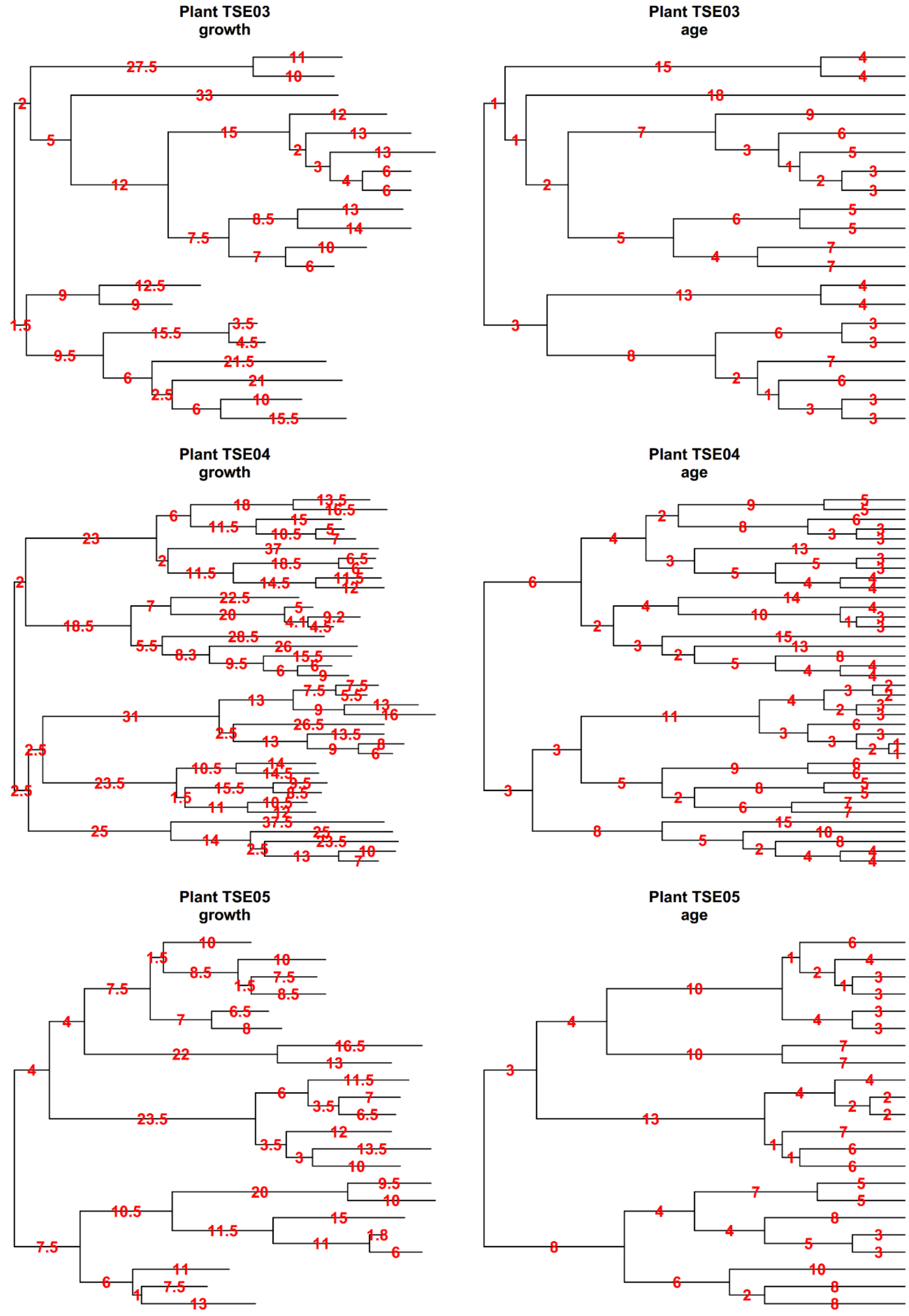

**Fig S2** Venn diagrams showing the distribution among the three *Lavandula latifolia* plants studied of the subindividually polymorphic methylation-sensitive AFLP markers.

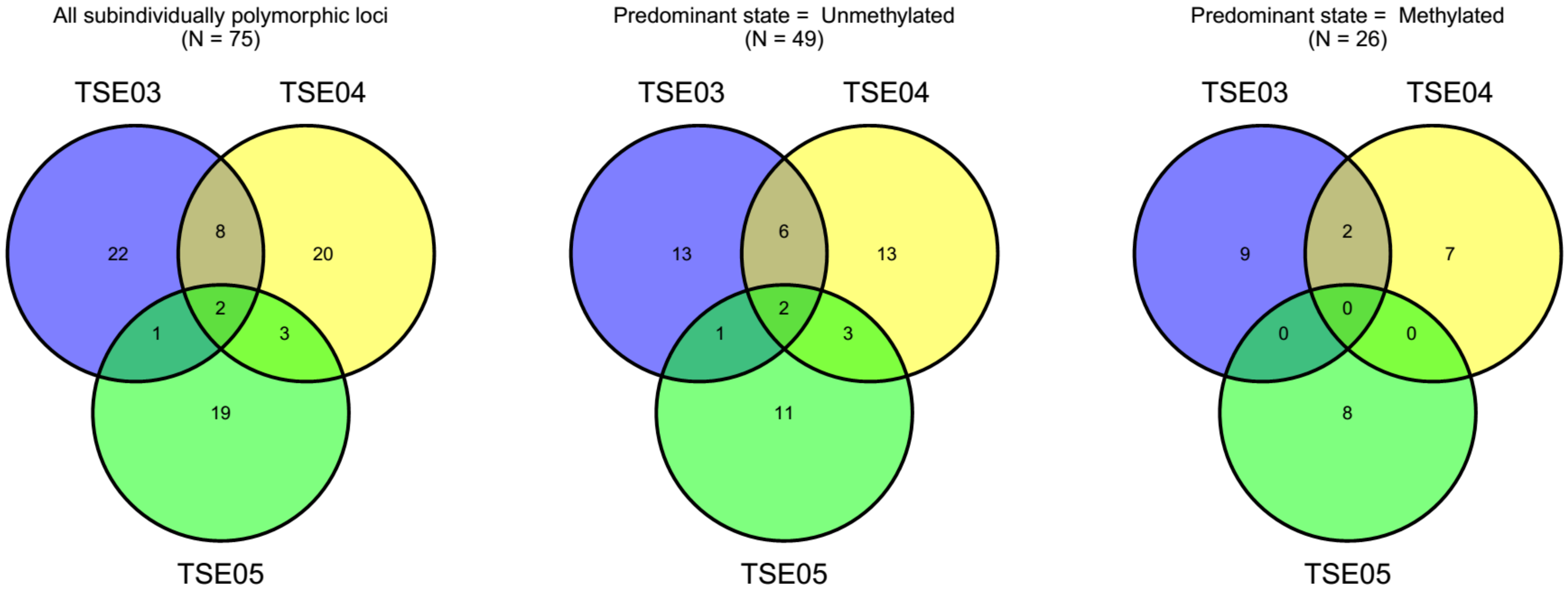

**Fig. S3** Within-plant genealogical character estimation of the methylation state of nine highly polymorphic MS-AFLP markers with significant genealogical signal. For each marker, methylation state in sampled modules (tree tips) and estimated posterior probabilities at nodes are coded as grey (methylated) or orange (unmethylated). Tree branch lengths represent linear distances between branching nodes and the units of horizontal axes are centimeters. Markers are identified by primer combination and fragment size in base pairs, and correspond to plants TSE03 (TG_CTA_297), TSE04 (TT_CAC_413, TA_CTA_278, TC_CGC_347, TA_CAC_251, TC_CCT_367) and TSE05 (TC_CGC_241, TC_CCT_200, TC_CCT_390).

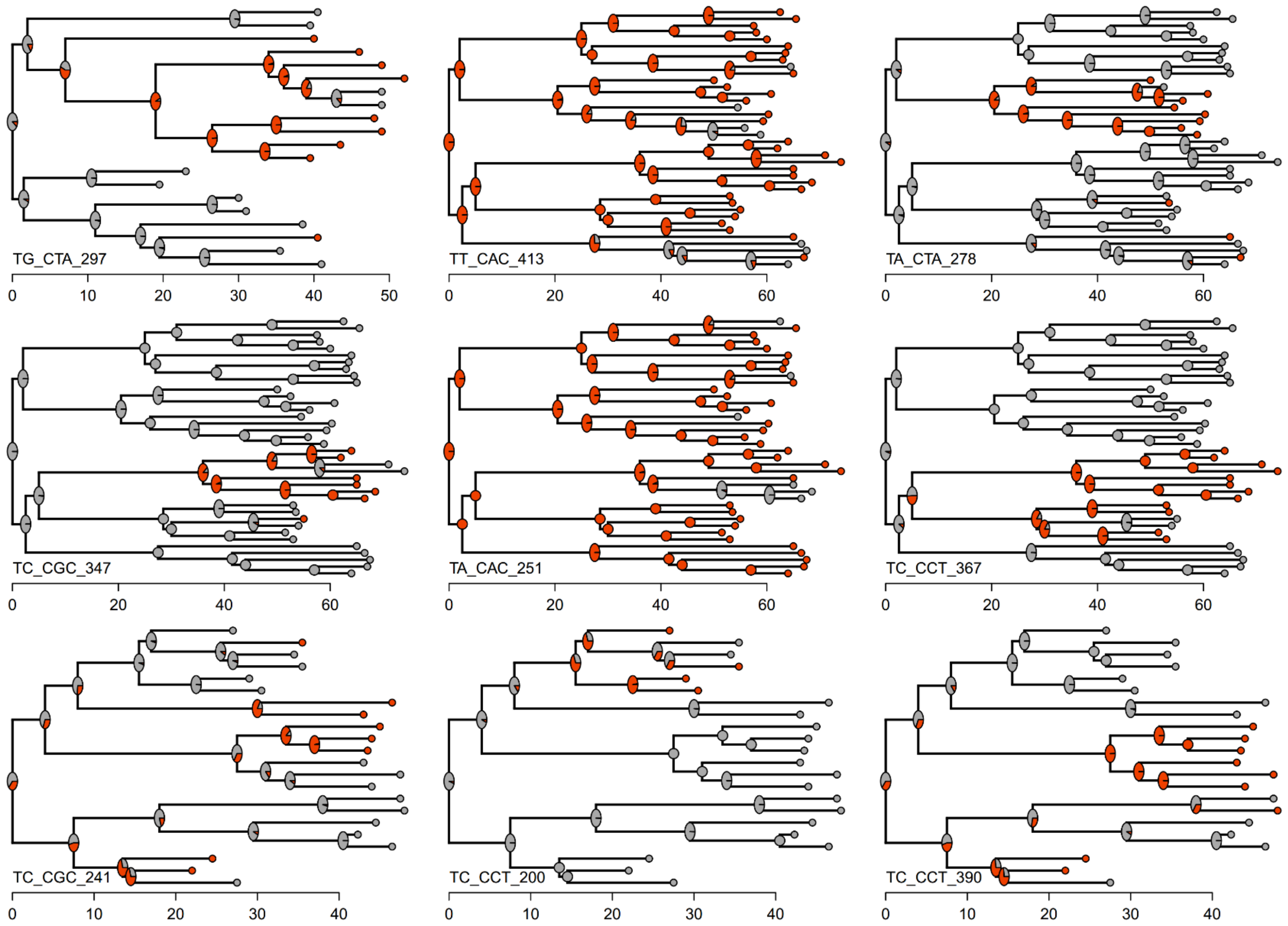

**Table S1.** Primer combinations and number of fragments in the size range 150-500 base pairs that were used in the methylation-sensitive, amplified fragment length polymorphism (MS-AFLP) analyses of DNA samples of leaves from 80 modules of three *Lavandula latifolia* plants.

| Primer combination | Number of fragments |
|---|---|
| *HpaII* TA– *Mse* CGT | 32 |
| *HpaII* TC– *Mse* CCT | 73 |
| *HpaII* TA– *Mse* CAC | 60 |
| *HpaII* TG– *Mse* CCT | 36 |
| *HpaII* TT– *Mse* CAC | 70 |
| *HpaII* TC– *Mse* CGC | 67 |
| *HpaII* TA– *Mse* CTA | 85 |
| *HpaII* TG– *Mse* CTA | 44 |
| All combined | 467 |

**Table S2** Summary of tests of within-plant genealogical signal, and fits of discrete character models (ER and ARD models), for highly polymorphic individual MS-AFLP markers.

| | | | Genealogical signal test (Fritz-Purvis *D* statistic) | | | Frequency of methylation states | | Parameter estimates from discrete character models [2] | | | | |
|---|---|---|---|---|---|---|---|---|---|---|---|---|
| Plant | MS_AFLP marker (HpaII_Mse) [1] | Tree branch metric | *D* statistic | $P$ ($D < 1$) | $P$ ($D \neq 0$) | Meth | Unmeth | ER.aic | ARD.aic | ER.q | ARD.q12 | ARD.q21 |
| TSE03 | TA_CTA_280 | Distance | 1.476 | 0.821 | 0.011 | 12 | 8 | 0.715 | 0.285 | 0.084 | 0.088 | 0.129 |
| | | Years | 1.456 | 0.832 | 0.006 | 12 | 8 | 0.698 | 0.302 | 2.837 | 2.255 | 3.383 |
| | TC_CGC_223 | Distance | -0.118 | 0.049 | 0.580 | 15 | 5 | 0.748 | 0.252 | 0.015 | 0.015 | 0.028 |
| | | Years | 0.136 | 0.086 | 0.451 | 15 | 5 | 0.735 | 0.265 | 0.029 | 0.041 | 0.115 |
| | TG_CTA_297 | Distance | -0.325 | 0.012 | 0.680 | 11 | 9 | 0.742 | 0.258 | 0.012 | 0.015 | 0.008 |
| | | Years | -0.228 | 0.012 | 0.633 | 11 | 9 | 0.739 | 0.261 | 0.031 | 0.038 | 0.014 |
| TSE04 | TA_CAC_251 | Distance | 0.097 | 0.020 | 0.464 | 6 | 32 | 0.728 | 0.272 | 0.005 | 0.018 | 0.006 |
| | | Years | 0.035 | 0.015 | 0.502 | 6 | 32 | 0.523 | 0.477 | 0.012 | 0.157 | 0.024 |
| | TA_CTA_278 | Distance | -0.365 | <0.0001 | 0.785 | 27 | 11 | 0.753 | 0.247 | 0.006 | 0.007 | 0.006 |
| | | Years | -0.423 | <0.0001 | 0.817 | 27 | 11 | 0.746 | 0.254 | 0.017 | 0.018 | 0.014 |

| | | | | | | | | | | | | |
|---|---|---|---|---|---|---|---|---|---|---|---|---|
| | TC_CCT_367 | Distance | -1.231 | <0.0001 | 0.998 | 26 | 12 | 0.538 | 0.462 | 0.003 | 0.000 | 0.012 |
| | | Years | -1.276 | <0.0001 | 0.998 | 26 | 12 | 0.438 | 0.562 | 0.006 | 0.000 | 0.028 |
| | TC_CGC_347 | Distance | -0.822 | <0.0001 | 0.913 | 31 | 7 | 0.653 | 0.347 | 0.004 | 0.003 | 0.010 |
| | | Years | -0.881 | <0.0001 | 0.931 | 31 | 7 | 0.381 | 0.619 | 0.009 | 0.000 | 0.078 |
| | TG_CCT_182 | Distance | 1.408 | 0.880 | 0.001 | 7 | 31 | 0.041 | 0.959 | 0.010 | 2.120 | 0.479 |
| | | Years | 1.463 | 0.915 | 0.001 | 7 | 31 | 0.031 | 0.969 | 0.025 | 0.720 | 0.164 |
| | TT_CAC_413 | Distance | 0.314 | 0.041 | 0.308 | 7 | 31 | 0.456 | 0.544 | 0.007 | 0.032 | 0.002 |
| | | Years | 0.250 | 0.031 | 0.348 | 7 | 31 | 0.572 | 0.428 | 0.018 | 0.079 | 0.007 |
| TSE05 | TC_CCT_200 | Distance | -0.400 | 0.029 | 0.646 | 18 | 4 | 0.149 | 0.851 | 0.009 | 0.000 | 0.053 |
| | | Years | -0.544 | 0.015 | 0.709 | 18 | 4 | 0.216 | 0.784 | 0.017 | 0.000 | 0.090 |
| | TC_CCT_390 | Distance | -0.688 | 0.001 | 0.862 | 13 | 9 | 0.566 | 0.434 | 0.014 | 0.001 | 0.023 |
| | | Years | -0.576 | <0.0001 | 0.835 | 13 | 9 | 0.705 | 0.295 | 0.025 | 0.027 | 0.005 |
| | TC_CGC_241 | Distance | -0.295 | 0.007 | 0.684 | 14 | 8 | 0.715 | 0.285 | 0.022 | 0.016 | 0.038 |
| | | Years | -0.082 | 0.010 | 0.571 | 14 | 8 | 0.756 | 0.244 | 0.037 | 0.033 | 0.014 |
| | TC_CGC_474 | Distance | 0.706 | 0.297 | 0.225 | 18 | 4 | 0.497 | 0.503 | 0.015 | 0.041 | 0.179 |
| | | Years | 0.737 | 0.287 | 0.203 | 18 | 4 | 0.355 | 0.645 | 0.026 | 0.080 | 0.348 |
| | TG_CTA_204 | Distance | 0.659 | 0.270 | 0.250 | 18 | 4 | 0.316 | 0.684 | 0.014 | 0.720 | 3.238 |

| | | | | | | | | | | | |
|---|---|---|---|---|---|---|---|---|---|---|---|
| | Years | 0.852 | 0.366 | 0.150 | 18 | 4 | 0.335 | 0.665 | 0.027 | 0.099 | 0.437 |
| TT_CAC_249 | Distance | 0.882 | 0.377 | 0.073 | 10 | 12 | 0.755 | 0.245 | 1.893 | 2.079 | 1.732 |
| | Years | 0.890 | 0.365 | 0.056 | 10 | 12 | 0.765 | 0.235 | 0.145 | 0.164 | 0.145 |

[1] Markers are identified by primer combination and fragment size (base pairs). See Supporting Information Table S1.

[2] ER.aic and ARD.aic: AIC weights for “equal rates” (ER) and “all rates different” (ARD) models, respectively. ER.q, estimate of methylation state transition rates under the ER model; ARD.q12 and ARD.q21, estimates of methylated-to-unmethylated and unmethylated-to-methylated transition rates under the ARD model.
